# Predictive Modeling of *Pseudomonas syringae* Virulence on Bean using Gradient Boosted Decision Trees

**DOI:** 10.1101/2021.05.27.445966

**Authors:** Renan N.D. Almeida, Michael Greenberg, Cedoljub Bundalovic-Torma, Alexandre Martel, Pauline W. Wang, Maggie A. Middleton, Syama Chatterton, Darrell Desveaux, David S. Guttman

## Abstract

*Pseudomonas syringae* is a genetically diverse bacterial species complex responsible for numerous agronomically important crop diseases. Individual *P. syringae* isolates are typically given pathovar designations based on their host of isolation and the associated disease symptoms, and these pathovar designations are often assumed to reflect host specificity although this assumption has rarely been rigorously tested. Here we developed a rapid seed infection assay to measure the virulence of 121 diverse *P. syringae* isolates on common bean (*Phaseolus vulgaris*). This collection includes *P. syringae* phylogroup 2 (PG2) bean isolates (pathovar *syringae*) that cause bacterial spot disease and *P. syringae* phylogroup 3 (PG3) bean isolates (pathovar *phaseolicola*) that cause the more serious halo blight disease. We found that bean isolates in general were significantly more virulent on bean than non-bean isolates and observed no significant virulence difference between the PG2 and PG3 bean isolates. However, when we compared virulence within PGs we found that PG3 bean isolates were significantly more virulent than PG3 non-bean isolates, while there was no significant difference in virulence between PG2 bean and non-bean isolates. These results indicate that PG3 strains have a higher level of host specificity than PG2 strains. We then employed machine learning to investigate if we could use genomic data to predict virulence on bean. We used gradient boosted decision trees to model the virulence using whole genome kmers, type III secreted effector kmers, and the presence/absence of type III effectors and phytotoxins. Our model performed best using whole genome data and was able to predict virulence with high accuracy (mean absolute error = 0.05). Finally, we functionally validated the model by predicting virulence for 16 strains and found that 15 (94%) had virulence levels within the bounds of estimated predictions. This study demonstrates the power of machine learning for predicting host specific adaptation and strengthens the hypothesis that *P. syringae* PG2 strains have evolved a different lifestyle than other *P. syringae* strains.

**AUTHOR SUMMARY:** *Pseudomonas syringae* is a genetically diverse Gammaproteobacterial species complex responsible for numerous agronomically important crop diseases. Strains in the *P. syringae* species complex are frequently categorized into pathovars depending on pathogenic characteristics such as host of isolation and disease symptoms. Common bean pathogens from *P. syringae* are known to cause two major diseases: the halo blight disease, which is characterized by large necrotic lesions surrounded by a chlorotic zone or halo of yellow tissue; and the bacterial spot disease, which is characterized by brown leaf spots. While halo blight can cause serious crop losses, bacterial spot disease is generally of minor agronomic concern. The application of statistical genetic and machine learning approaches to genomic data has greatly increased our power to identify genes underlying traits of interest, such as host specificity. Machine learning models can be used to predict outcomes from new samples or to identify the genetic feature(s) that carry the most importance when predicting a particular phenotype. Here, we implemented a rapid method for screening a proxy of virulence for *P. syringae* isolates on common bean, and used this screen to assess virulence of *P. syringae* strains on bean. We found that halo blight pathogens display a stronger degree of host specificity compared to brown spot pathogens, and that genomic kmers and virulence factors can be used to predict the virulence of *P. syringae* isolates on bean using machine learning models.

## INTRODUCTION

*Pseudomonas syringae* is a genetically diverse Gammaproteobacterial species complex responsible for numerous agronomically important crop diseases [1–4]. Strains in the *P. syringae* species complex are frequently categorized into pathovars depending on pathogenic characteristics such as host of isolation and disease symptoms [5, 6]. The species complex is also subdivided into phylogenetic groups (i.e., phylogroups, PGs) based on multilocus sequence typing or genomic analysis [1, 7–10]. Currently, there are 13 recognized PGs [7], of which seven have been termed primary PGs based on their genetic relatedness and the near universal presence of the canonical *P. syringae* type III secretion system (discussed below) [1, 9].

Phylogenetic analyses of *P. syringae* isolates suggest that adaptation to specific hosts has evolved multiple times in the evolutionary history of the species complex. For example, cherry and plum pathogens are found in clades distributed in PG1, PG2, and PG3 [11, 12], hazelnut pathogens are distributed among two distinct clades in PG1 and PG2 [13], while pathogens of common bean (including snap, green, kidney, and French bean) are found in PG2 and PG3 [9].

Common bean pathogens from *P. syringae* PG3 are generally classified as pathovar *phaseolicola* and are responsible for halo blight disease, which is characterized by large necrotic lesions surrounded by a chlorotic zone or halo of yellow tissue [14–16]. Bean pathogens of PG2 are generally classified as pathovar *syringae* and are responsible for bacterial spot disease, which is characterized by brown leaf spots [17, 18]. While halo blight can cause serious crop losses, bacterial spot disease is generally of minor agronomic concern. The PG3 *phaseolicola* bean isolates show a high degree of phylogenetic clustering, with most strains sharing a relatively recent common ancestor that is closely related to a compact sister clade of soybean pathogens [9]. In contrast, PG2 *syringae* bean isolates show very little phylogenetic clustering and are frequently more closely related to non-bean isolates than other bean isolates [9].

Assuming that host specificity is a heritable trait, the exploitation of a common host by divergent lineages of strains can be explained by several different mechanisms, including: 1) evolution via shared, vertically transmitted host specificity factors; 2) convergent evolution via unrelated genetic mechanisms; or 3) convergent evolution via the horizontal acquisition of host specificity factors from divergent lineages. Another layer of complexity is that host specificity could come about either through the gain of genetic factors that promotes growth on a new host, or alternatively, by the loss of a factor that otherwise limits growth (e.g., by inducing a host immune response). In fact, the most thorough study of host convergence in *P. syringae* suggests that isolates can make use of multiple mechanisms simultaneously [11, 12]. For example, diverse lineages of cherry pathogens have exchanged and lost key genes and used multiple mechanisms to successfully infect this host [11, 12].

One of the most important and dynamic classes of *P. syringae* virulence and host specificity factors are type III secreted effectors (T3SEs). T3SEs are proteins translocated through the type III secretion system directly into the eukaryotic host cell where they interfere with host immunity or disrupt cellular homeostasis to promote the disease process. There are 70 distinct families of *P. syringae* T3SEs, and most strains carry a suite of T3SEs consisting of 12 to 50 T3SEs, with an average of ∼30 [19]. Plants have responded to T3SEs by evolving immune receptors and complexes that trigger an effector-triggered immune (ETI) response when they detect the presence or activity of a T3SE [20, 21]. Consequently, the outcome of any particular host-microbe interaction depends to a large degree on the specific T3SE profile of the pathogen and the complement of immune receptors carried by the host. The strong selective pressures imposed by the host-microbe arms race results in dynamic evolution of T3SEs in general, with frequent horizontal transmission, acquisition, and loss [9, 19, 22].

The suites of T3SEs carried by PG2 and PG3 strains vary in size, with PG2 strains carrying an average of ∼19 T3SEs vs. ∼27 for PG3 strains [19]. PG2 strains are also known to carry more phytotoxins, which contribute to virulence and niche competition via a variety of mechanisms such as membrane disruption and hormone mimicry [3, 9, 23]. In general, PG2 strains are believed to show lower levels of host specificity and better ability to survive on leaf surfaces (i.e., epiphytic growth) [1, 3, 24, 25].

The application of statistical genetic and machine learning approaches to genomic data has greatly increased our power to identify genes underlying traits of interest, such as host specificity [26]. Statistical genetic approaches like genome-wide association studies (GWAS) are well developed for studying human traits and have more recently gained traction in the study of bacterial traits as statistical and phylogenetic methods have been developed to handle the shared evolutionary history of segregating genetic variants (i.e., population structure) [27–32]. While GWAS approaches have great power for finding genotype-phenotype associations, they generally measure associations on a locus-by-locus basis, and therefore can miss more complex interactions among loci that impact traits. An alternative approach for predicting genotype-phenotype associations is to use machine learning, which generally describes a large range of statistical approaches that create models derived from a dataset consisting of features (e.g., genetic variants) linked to a trait or outcome (e.g., host specificity). These models can be used to predict outcomes from new samples or to identify the feature(s) that carry the most importance in the model. Although machine learning approaches are better suited for identifying suites of interacting genetic variants than GWAS, they are more limited in their ability to deal with complex evolutionary relationships among these variants [33, 34].

Here, we implemented a rapid method for screening a proxy of virulence for *P. syringae* isolates on common bean. We used this screen to assess virulence of 121 strains from nine PGs on bean, and then expand the strain collection by imputing the virulence for an additional set of isolates based on their core genome relationship to the screened strains. While most of bean pathogens come from PG2 and PG3, we found that PG3 pathogens display a stronger degree of host specificity compared to PG2 pathogens. Finally, we developed a random forest regression model using genomic kmers and virulence factors as features to predict the virulence of *P. syringae* isolates on bean.

## RESULTS

### Genome analysis

We characterized the genomic diversity of the isolates used in this study via phylogenetic analysis (Fig. 1) and by measuring the pairwise core genome synonymous substitution rates (Ks) and pairwise accessory genome Jaccard distances for each isolate combination. The 36 bean halo blight pathogens of pathovar *phaseolicola* largely cluster in one very closely related clade in PG3 and had a core genome Ks of 0.0039 and an accessory genome Jaccard distance of 0.28 compared to the entirety of 142 PG3 strains, which had a core genome Ks of 0.047 and an accessory genome Jaccard distance of 0.23. In contrast the 21 bean spot disease pathogens of pathovar *syringae* are broadly distributed throughout PG2 and had a core genome Ks of 0.1218 and an accessory genome Jaccard distance of 0.51 compared to the entirety of 66 PG2 strains, which had a core genome Ks of 0.1223 and an accessory genome Jaccard distance of 0.27 (Fig. 2). Due to their clonal nature, PG3 bean isolates were found to have a considerably higher number of gene families in the hard (present in 100% of the isolates) and soft core (present in >95% of the isolates) genomes, as well as a lower number of singleton families (present in a single isolate) in comparison to PG 2 bean isolates, despite the higher number of PG3 samples analyzed (Fig. 3).

**Figure 1:**
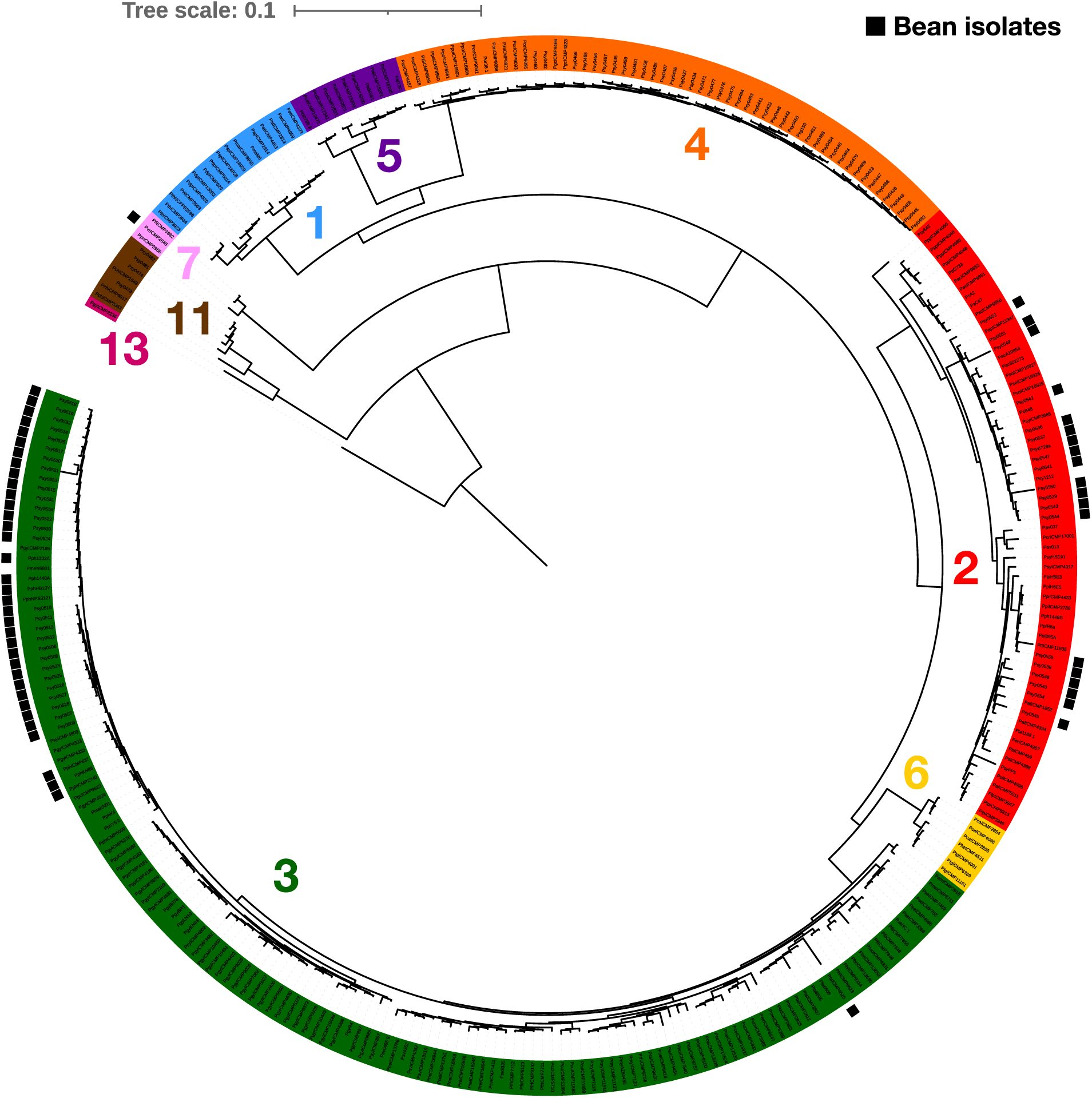
Distribution of bean isolates across the *Pseudomonas syringae* core genome phylogeny. Core genome phylogeny of 320 *P. syringae* isolates. Clade numbers represent phylogroups. Metadata row 1 shows strains isolated from diseased bean plants.

**Figure 2:**
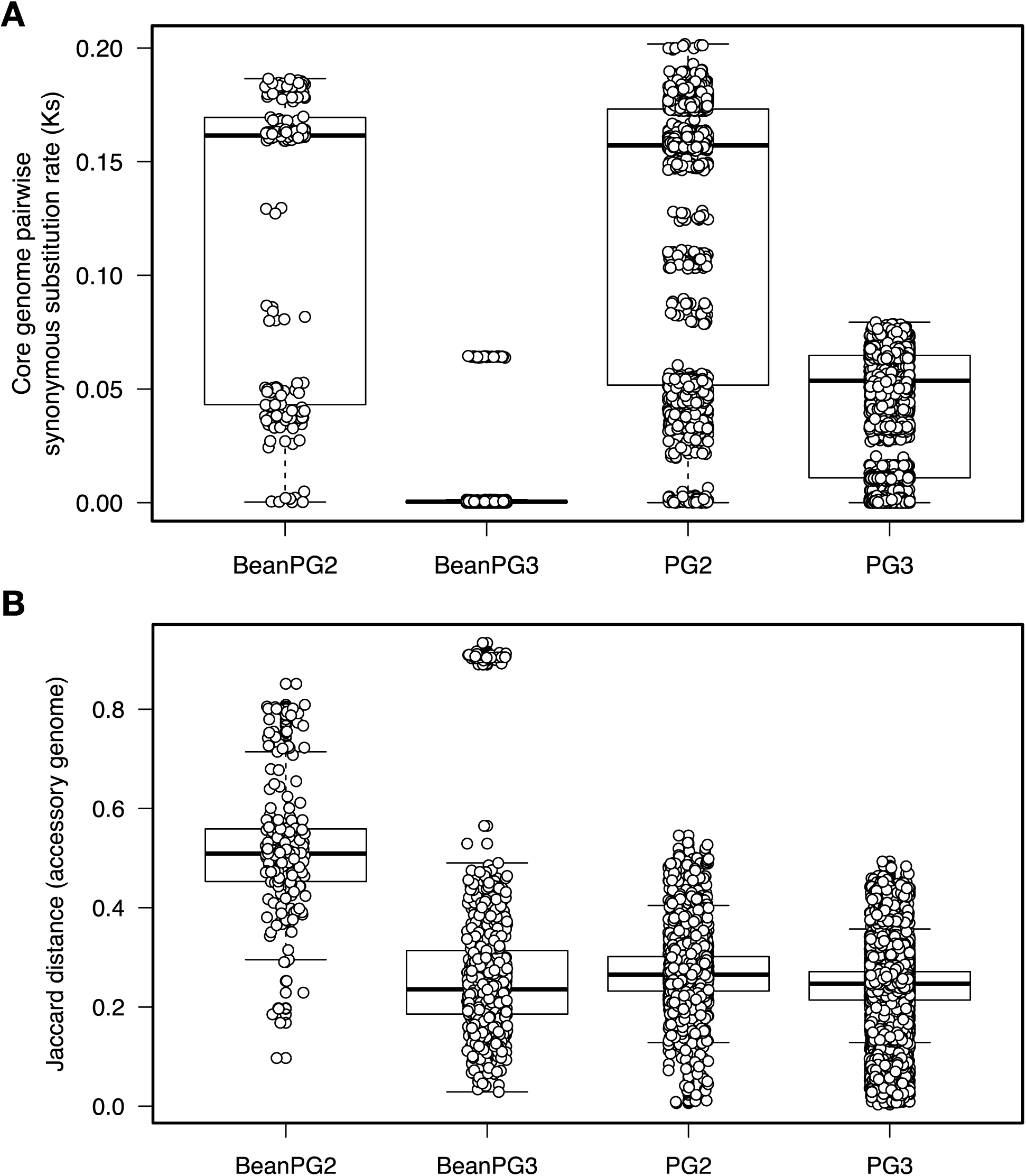
Core and accessory genome diversity. Comparison of (A) core genome synonymous substitution rate (Ks) and (B) accessory genome Jaccard distance for 21 PG2 bean isolates, 36 PG3 bean isolates, and all 66 isolates from PG2 and 142 isolates from PG3.

**Figure 3:**
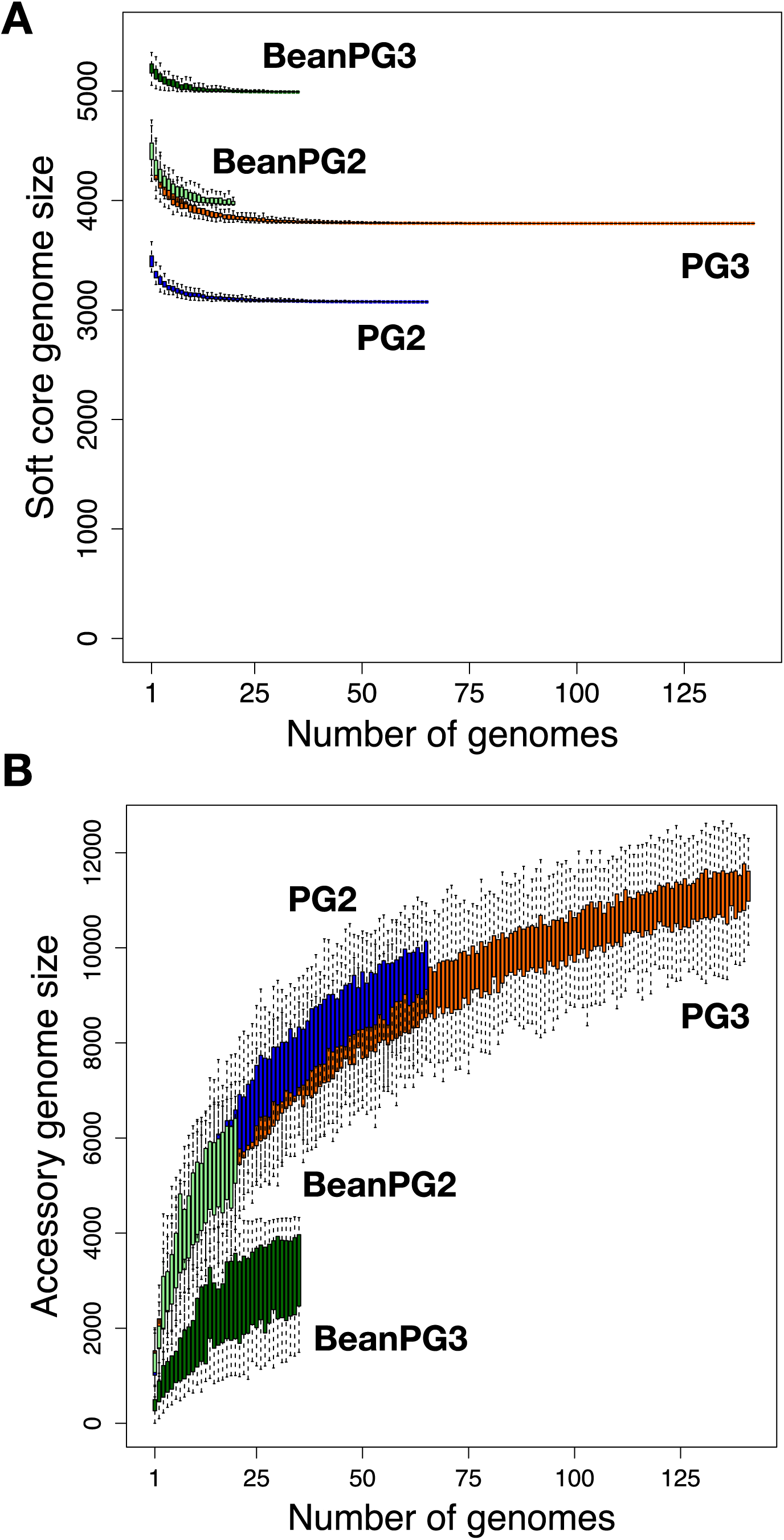
Rarefaction curves for the core and accessory genomes. Families present in 95% (soft core genome) of *P. syringae* isolates exponentially decay as each new genome is added to the analysis. The total number of gene families identified continues to increase indefinitely with the addition of new genomes when singletons (families only present in one isolate) are included.

### Virulence screen development

We developed a high-throughput seed infection assay to measure the virulence of *P. syringae* isolates on common bean. We used seed infection based on the understanding that many bean infections are caused by contaminated seeds [14, 45–47]. These infections can reduce overall plant health, which is reflected in plant fresh weight. For the screen, we soaked bean seeds (*P. vulgaris* var. Canadian Red) in a *P. syringae* suspension (∼5×10^5^ cells / ml) for 24 hours prior to planting. Plant fresh weight was then determined after 14 days. We first assessed if plant weight was correlated with *in planta* bacterial load by comparing our seed infection assay to the traditional syringe infiltration virulence assay using 24 *P. syringae* isolates from 9 out of the 13 PGs (Fig. 4). Well-established bean pathogens such as PG3 strain *P. syringae* pv. *phaseolicola* 1448A (Pph1448A) [14, 16] and the PG2 strains *P. syringae* pv. *syringae* B728a (PsyB728a) [17, 48] showed the highest levels of bacterial growth and lowest plant weights, while the other isolates from PGs 1-7, 11, and 13 showed a range of values. Overall, there was a significant negative association between bacterial growth and plant weight (R^2^=0.63, P+5.0e-6), supporting the use of seed infection and plant fresh weight to assess bacterial virulence.

**Figure 4:**
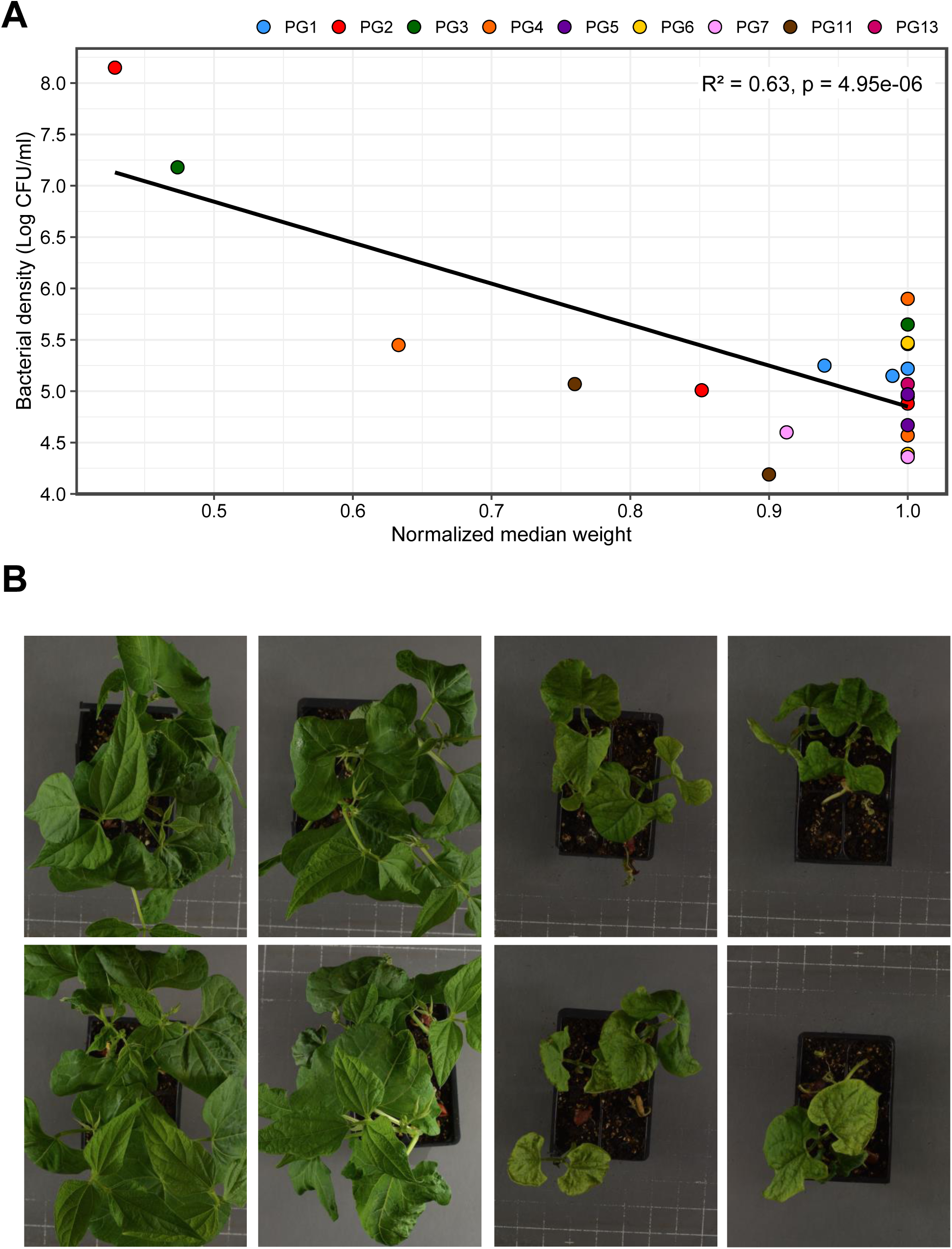
Correlation between bacterial load following pressure infiltration of mature bean leaves and plant weight following seed infection. (A) Bacterial density in 14 day old infected bean leaves (pressure infiltration 3 days post infection) as a function of normalized median weight of seed infected plants at 14 days old. There is a strong negative correlation between bacterial density and plant weight across 24 *P. syringae* isolates (linear regression; *F* = 36.95, df = 21, *p* = 4.95e-06, R^2^ = 0.62). (B) Characteristic plant phenotypes of 14 day old plants following seedling infection. The photos on the left are for plants infected with MgSO_4_, while the photos on the right shows plants infected with the bean pathogen Psy0540.

To determine the power of this assay, we performed initial seed infection trials with six *P. syringae* isolates and 50 or more replicate plants. We used a rarefaction analysis of normalized plant weights to determine the number of replicate plants required to distinguish pathogens from non-pathogens with >95% confidence (Tukey-HSD test) and found that the test power plateaued at ∼20 replicates per treatment (Fig. S1). Therefore, we performed future seed infection assays using 30 replicate plants per treatment.

### Virulence screen

We screened 121 non-clonal representative *P. syringae* isolates from nine PGs to assess the virulence potential as measured by reduced plant fresh weight in 14-day old bean plants after seed infection (Fig. 5). This screened set was subset of the non-clonal set selected to maximize coverage of the species complex while focusing on a manageable number for screening. The screened strains resulted in a highly skewed distribution of normalized fresh weights (i.e., virulence), with a mean of 0.78, median of 0.94, and standard deviation of 0.30. An examination of the 30 strains in the first quartile revealed normalized fresh weights between 0.00 and 0.61, with 56.7% (17 strains) being bean isolates (Fig. 6). These 17 strains represent 58.6% of all 29 bean isolates screened.

**Figure 5:**
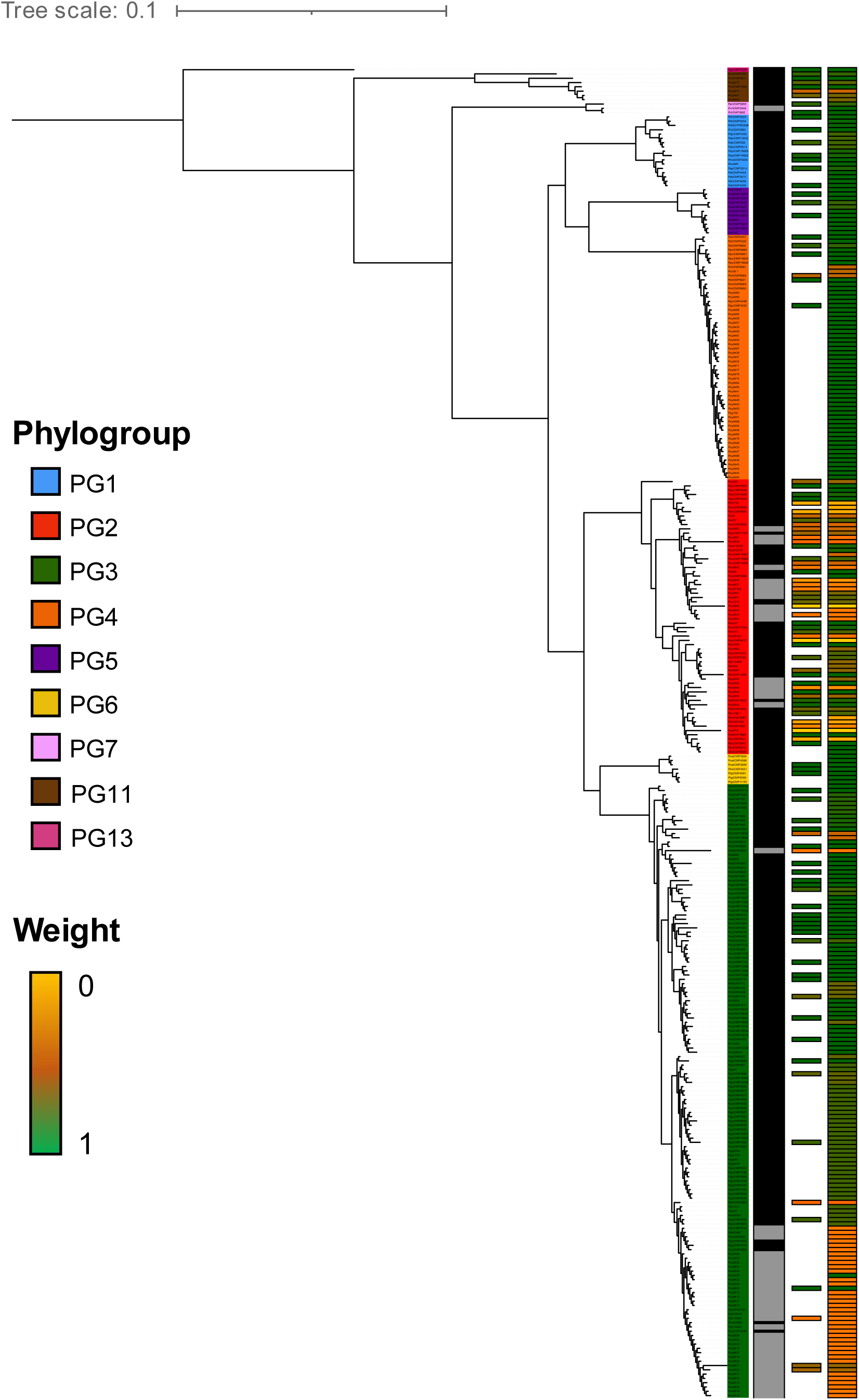
Core genome phylogeny of 320 *P. syringae* isolates. Leaf colors represent PG affiliation as shown in the legend. Metadata row 1 shows bean isolates in black and non-bean isolates in gray. Metadata row 2 shows normalized median bean weight after seed infection by the respective isolates. Green indicates a high weight, while yellow indicates a low weight as show in the legend. Metadata row 3 shows normalized median bean weight for the expanded strain collection based on phylogenetic imputation.

**Figure 6:**
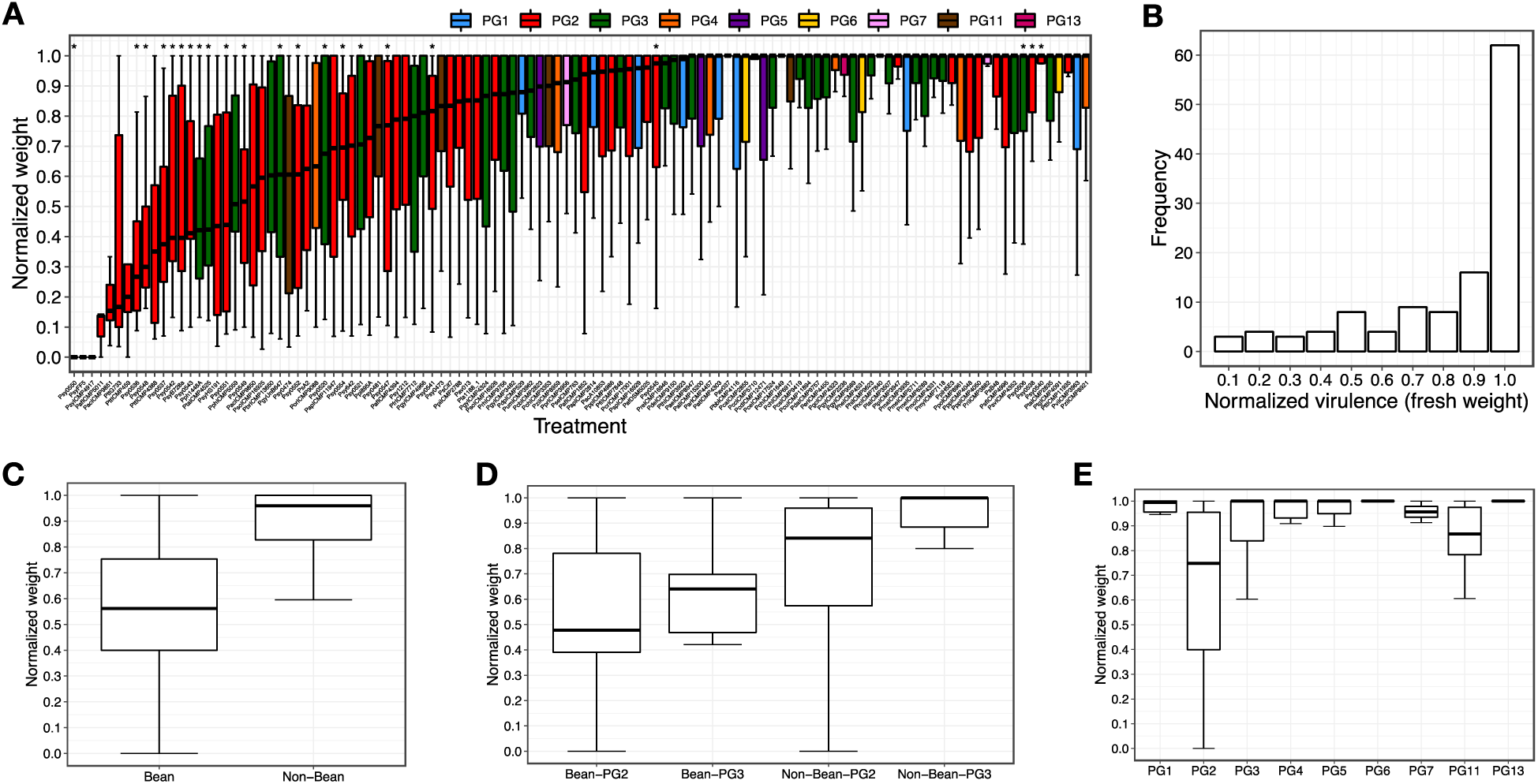
Virulence stratified by host and phylogroup. (A) Boxplots showing the distribution of virulence (i.e., normalized plant weight 14 days after seed infection) ordered by the median virulence. Colors correspond to phylogroups as shown along the top. Bean isolates are indicated with asterisks above their respective boxplots. (B) Frequency plot of virulence for the 121 screened *P. syringae* strains. Distribution of virulence values for (C) bean verses non-bean isolates, (D) bean and non-bean isolates stratified by phylogroup, and (E) virulence values stratified by phylogroup (PG) for the set of 121 screened strains.

Significant differences in virulence, as measured by normalized fresh weight, were observed when comparing the strain collection stratified by host of isolation and PG (Table 1). The 29 bean isolates had an average virulence (normalized fresh weight) of 0.59 compared to 0.85 for the 92 non-bean isolates (p=4.2e-5, 2-tailed, heteroscedastic T-test, same for tests discussed below). As all the bean isolates are found in PG2 and PG3, we compared the virulence of bean isolates to non-bean isolates within these two PGs individually and found no significant difference for PG2 (p=0.128) but a strong difference for PG3 (p=8.9e-4). Additionally, there were no significant differences between PG2 and PG3 bean isolates (p=0.460). We then looked for differences in virulence between strains from different PGs irrespective of their host of isolation (only comparing PGs with at least six tested strains, using 2-tailed, heteroscedastic T-tests, Bonferroni corrected for seven total tests), and found that strains in PG2 were significantly more virulent on bean than strains from PG1, PG3, and PG4 (p= 1.33e-07, 0.012, and 0.029 respectively), but not relative to PG6. In contrast, strains from PG3 were only significantly more virulent on bean than strains from PG1 (p=0.006). No other significant pairwise PG comparisons were observed. Interestingly, we noticed that PG2 non-bean isolates showed higher virulence on bean than non-bean isolates from other PGs (p=2.87e-4). This indicates that PG2 isolates show greater virulence on bean *irrespective* of host of isolation, although the degree of virulence is relatively low. This pattern was reversed in PG3 where non-bean isolates had significantly lower virulence than non-bean isolates from all other PGs (p=4.52e-3).

**Table 1.** Virulence and Germination Assay Summary

|  | N | Virulence <sup>2</sup> |  |  | Germination Frequency (%) |  |  |
| --- | --- | --- | --- | --- | --- | --- | --- |
| Group <sup>1</sup> | Strains | Mean | Median | Stdv | Mean | Median | Stdv |
| All Strains | 121 | 0.789 | 0.913 | 0.271 | 60.10 | 63.33 | 24.31 |
| PG1 | 8 | 0.972 | 0.995 | 0.043 | 72.99 | 79.17 | 15.71 |
| PG2 | 50 | 0.658 | 0.748 | 0.320 | 46.74 | 53.33 | 26.41 |
| PG3 | 42 | 0.841 | 0.964 | 0.218 | 68.15 | 67.50 | 16.53 |
| PG4 | 6 | 0.924 | 1.000 | 0.147 | 83.61 | 82.50 | 4.52 |
| PG5 | 3 | 0.966 | 1.000 | 0.059 | 73.33 | 78.33 | 10.14 |
| PG6 | 3 | 1.000 | 1.000 | 0.000 | 80.56 | 81.67 | 3.47 |
| PG7 | 2 | 0.956 | 0.956 | 0.062 | 85.00 | 85.00 | 7.07 |
| PG11 | 6 | 0.851 | 0.867 | 0.151 | 43.89 | 50.00 | 20.75 |
| PG13 | 1 | 1.000 | 1.000 | NA | 91.67 | 91.67 | NA |
| All Bean Isolates | 29 | 0.594 | 0.516 | 0.271 | 45.89 | 53.33 | 23.52 |
| All Non-Bean | 92 | 0.850 | 0.960 | 0.244 | 64.58 | 68.33 | 23.05 |
| PG2 Bean | 16 | 0.560 | 0.478 | 0.293 | 36.84 | 32.22 | 26.42 |
| PG2 Non-Bean | 34 | 0.704 | 0.841 | 0.327 | 51.41 | 56.67 | 25.47 |
| PG3 Bean | 13 | 0.635 | 0.606 | 0.246 | 57.02 | 58.89 | 13.21 |
| PG3 Non-Bean | 29 | 0.933 | 1.000 | 0.122 | 73.14 | 73.33 | 15.56 |

<sup>1</sup> PG = phylogroup
<sup>2</sup> Normalized fresh weight 2 weeks after seed infection

### Germination screen

We then assessed whether the virulence of a strain also influenced the germination frequency of bean seeds. Pathogenic microbes are known to interfere with the seed germination both through the direct action of phytotoxins and the indirect action of immune activation [23, 45, 46, 49, 50]. In fact, seedling growth inhibition is a well-established assay for immune activation in *Arabidopsis thaliana* [50]. In general, the frequency distribution for bean germination inhibition was less skewed than the frequency distribution for virulence, with mean = 60.1%, median = 63.3%, and standard deviation = 24.3% (Fig 7, Table 1). The average germination frequency for all bean isolates was 45.9% compared to 64.6% for non-bean isolates (p=4.92e-4). When stratifying the bean isolates by PG, we found no significant difference in germination frequency between PG2 bean and non-bean isolates (p=0.076), while the comparison was significant for PG3 (p=0.002). However, in contrast to the virulence assays we observed a slightly significant difference between germination frequency for PG2 bean isolates and PG3 bean isolates (p=0.014). Other inter-phylogroup comparisons were similar to what was found for the virulence assays, PG2 strains were significantly different from strains from PG1, PG3, and PG4 (p=0.010, 6.23e-5, 8.53e-11 respectively 2-tailed, heteroscedastic T-test, Bonferroni corrected for seven tests), while PG3 strain were also significantly different from PG4 (p=2.23e-4). Also similar to the virulence results, non-bean PG2 isolates resulted in a lower germination frequency than all other non-bean isolates (p=9.60e-5), while non-bean PG3 strains had a higher germination frequency than non-bean isolates from all other PGs (p=0.004), although this pattern is absent when non-bean PG2 strains were removed from the analysis.

**Figure 7:**
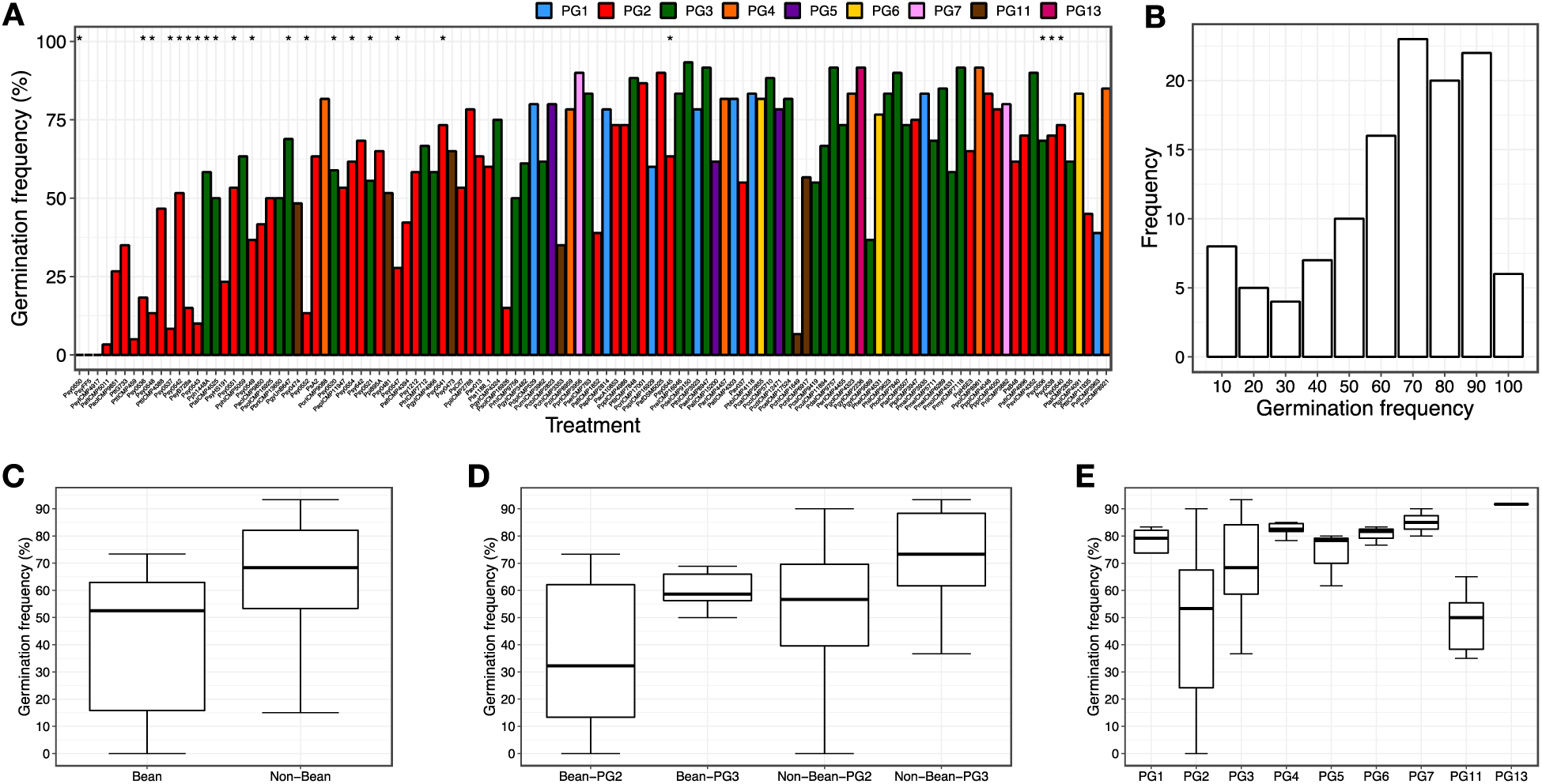
Germination frequencies by host and phylogroup. (A) Boxplots showing the germination frequencies frequency for 121 screened *P. syringae* strains with strains presented in the same manner and order as in Fig. 6. (B) Frequency plot of germination for the 121 screened strains. Distribution of germination frequencies for (C) bean verses non-bean isolates, (D) bean and non-bean isolates stratified by phylogroup, and (E) virulence values stratified by phylogroup (PG) for the set of 121 screened strains.

Finally, we measured the association between virulence (i.e., normalized fresh weight) and germination frequency and found a strong association between the two metrics for the full dataset (R^2^=0.46, p=2.2e-16). Stratifying by PG and bean isolates showed a strong association for PG2 bean isolates (linear regression; *F* = 28.91, df = 13, p=0.0001, R^2^ = 0.68), but no significant association for PG3 bean isolates (R^2^=0.62, p=0.06) (Fig 8).

**Figure 8:**
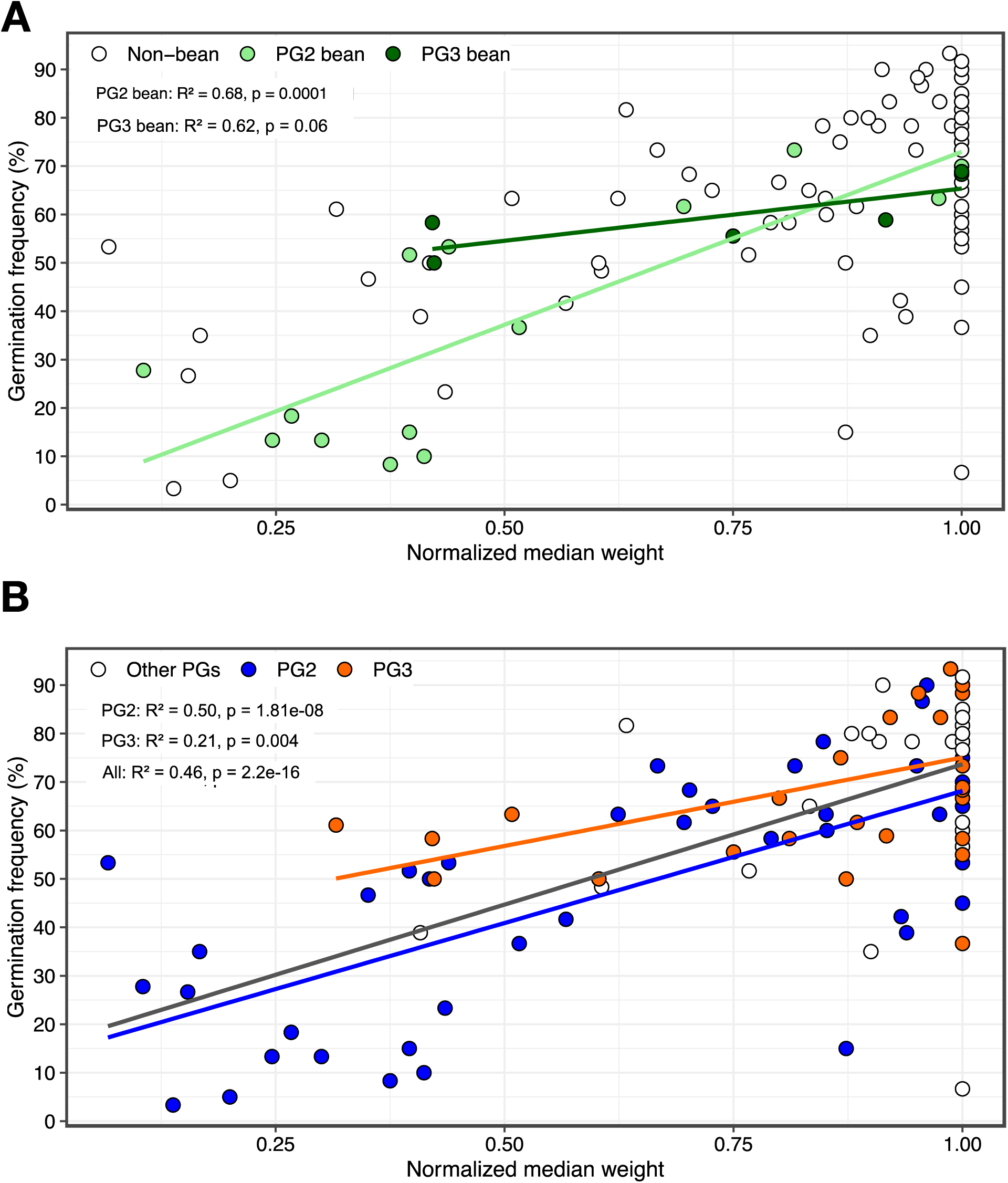
Correlation between germination frequency and virulence of seed infected bean plants. (A) Germination frequency vs virulence (i.e., normalized fresh weight) stratified by PG2 and PG3 bean isolates. A strong correlation is found between germination frequency and virulence of bean strains from PG2 (linear regression; *F* = 28.91, df = 13, *p* = 0.00012, r^2^ = 0.66). (B) Germination frequency vs. virulence stratified by PG irrespective of host.

### Predictive modelling of *P. syringae* virulence on bean

We used a gradient boosted decision tree regression model to predict the virulence of *P. syringae* isolates on beans. The model used one of three input feature classes: 1) genomic kmers; 2) T3SE kmers; or 3) presence / absence of T3SEs and phytotoxins. T3SEs and phytotoxins are well-known virulence factors, with the former often strongly associated with host specificity. Plant weight 14 days after seed infection was used as the continuous outcome variable in our model. We could have also used seed germination frequency in this assay but felt that plant fresh weight more accurately reflected the virulence concerns of bean producers. The goal of analysis was to assess the power of machine learning to predict disease outcomes based on genome sequences and to predict the host specificity of new isolates based on their genome sequence.

We used two nested collections of strains to generate the model. The first collection was comprised of the 121 of the isolates directly screened for virulence, which was made of 29 bean isolates and 92 non-bean isolates, including 50 PG2 isolates (16 bean, 34 non-bean), and 42 PG3 isolates (13 bean, 29 non-bean). This collection is slightly smaller than the full screened set since it does not include the additional PG3 bean isolates added to balance the experimental design (q.v., materials and methods). The second collection was an expanded strain set in which we imputed virulence values based on genomic similarity. The imputation process involved identifying strains in our collection belonging to the same clonal lineage as those assayed in our virulence screen (i.e., having a core genome evolutionary distance of less than 0.001 and a T3SE Jaccard similarity of greater than 0.8 to a screened strain). Any strains meeting these criteria were assigned the same virulence as the corresponding screen strain. This imputation process almost tripled the size of our sample set, resulting in an expanded collection of 320 strains, which was made of 59 bean isolates and 261 non-bean isolates, including 66 PG2 isolates (19 bean, 47 non-bean), and 142 PG3 isolates (39 bean, 104 non-bean). We also trained a model on PG2 and PG3 strains separately since bean isolates from these PGs interact with their host very differently.

Our analysis revealed a significant linear relationship between sample size and the overall performance of the models (Fig. 9, MAE linear regression *F* = 5.89, df = 10, p=0.03, R^2^ = 0.3). Hence, the data sets with the least number of samples consistently showed the poorest performance regardless of the input data (e.g., PG2 and the screened collection). Furthermore, our models achieved the highest predictive power when trained exclusively at isolates from PG3, which may be explained by the strong clonal separation of bean vs. non-bean pathogens within this PG which results in the statistical reinforcement of features with improved predictive power during model training. Perhaps not surprisingly, models built using the greatest diversity, with respect to number of strains (i.e., the expanded strain collection) and number of genetic feature (i.e., whole genome kmers) showed the highest predictive power, with the best model having a mean absolute error (MAE) of 0.06 (Table 2).

**Figure 9:**
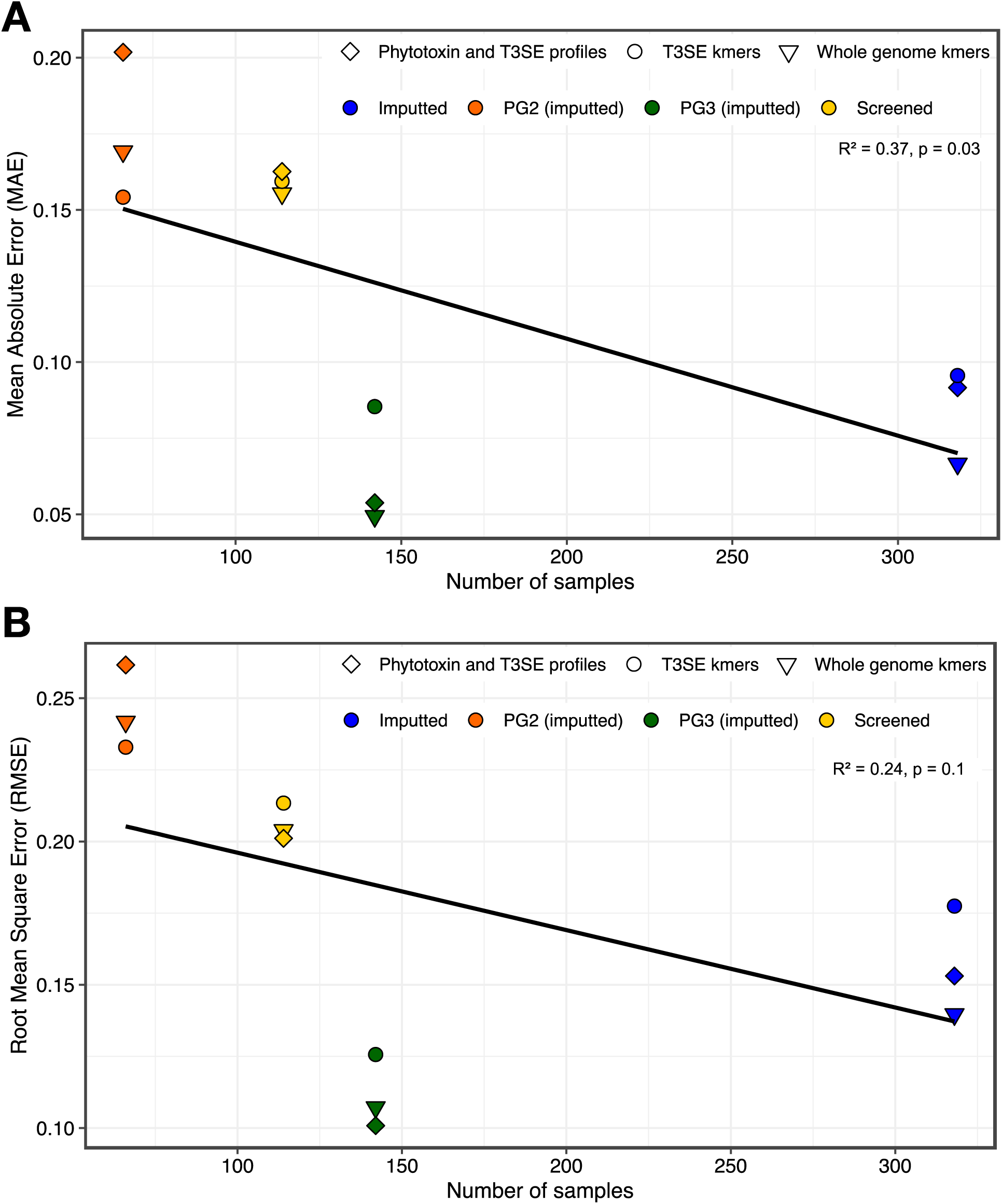
Performance of supervised machine learning models on virulence predictions with genome data. Models trained with the screened collection (yellow), imputed expanded collection (blue), imputed expanded PG2 set (orange), and imputed expanded PG3 set (green). Diamonds, circles, and triangles represent models trained with a phytotoxin and T3SE binary matrix, T3SE kmers, and whole genome kmers, respectively. (A) Mean Absolute Error (MAE) as a function of number of samples across 50 cross-validation splits. Increasing the number of samples seen by the model significantly increases model performance (linear regression; *F* = 5.89, df = 10, *p* = 0.03, r^2^ = 0.37). (B) Root Mean Square Error (RMSE) as a function of number of samples across 50 cross-validation splits. Models trained on whole genome kmers with a higher number of samples consistently outperform other models.

**Table 2.** Machine Learning Model Performance

| Group | Samples | Dataset | MAE | RMSE |
| --- | --- | --- | --- | --- |
| PG3 | 142 | Whole genome kmers | 0.049 | 0.107 |
| PG3 | 142 | Toxin and T3SE binary matrix | 0.054 | 0.101 |
| Imputed | 320 | Whole genome kmers | 0.067 | 0.140 |
| PG3 | 142 | T3SE kmers | 0.085 | 0.126 |
| Imputed | 320 | Toxin and T3SE binary matrix | 0.092 | 0.153 |
| Imputed | 320 | T3SE kmers | 0.096 | 0.177 |
| PG2 | 66 | T3SE kmers | 0.154 | 0.233 |
| Screened | 121 | Whole genome kmers | 0.155 | 0.204 |
| Screened | 121 | T3SE kmers | 0.159 | 0.213 |
| Screened | 121 | Toxin and T3SE binary matrix | 0.163 | 0.201 |
| PG2 | 66 | Whole genome kmers | 0.169 | 0.242 |
| PG2 | 66 | Toxin and T3SE binary matrix | 0.202 | 0.262 |

### Model functional validation

Finally, we evaluated the power of our gradient boosted decision tree regression model on unseen strains by using the kmer profiles of *P. syringae* isolates not previously examined to predict virulence and comparing these predictions to actual virulence measures obtained through the seed infection virulence assay. Importantly, none of the strains in the functional validation set were clonally related to any screened strain, i.e., having a core genome evolutionary distance >0.001 and a T3SE Jaccard similarity of <0.8 compared to the screened strains. When comparing actual to predicted virulence of the 16 strains in the functional validation set, we found that 15 (94%) strains had virulence levels within the bounds of estimated predictions given the calculated RMSE values (±0.20) (Fig. 10).

**Figure 10:**
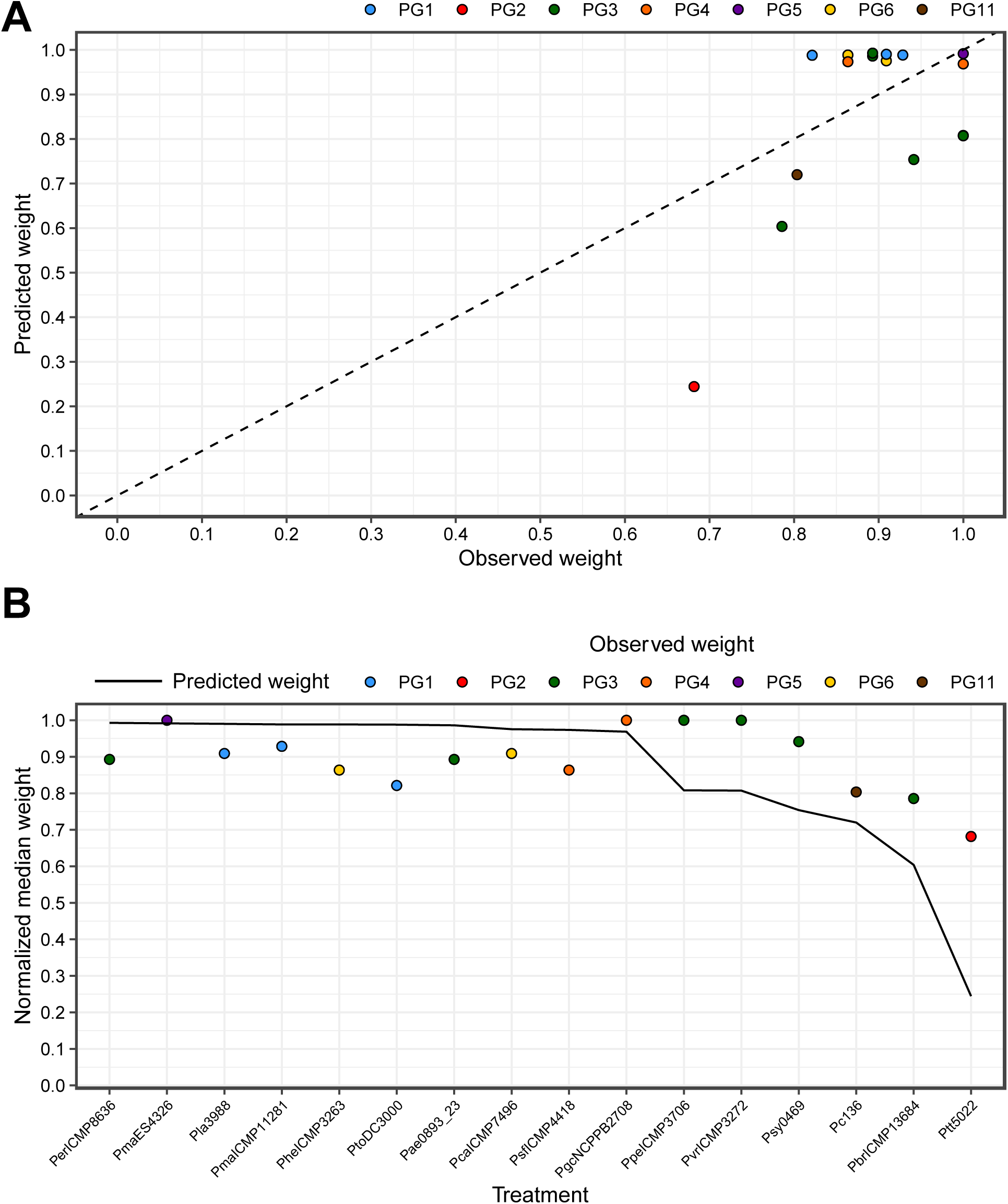
Correlation between predicted and observed weights of plants infected with isolates previously unseen by the model. (A) Predicted virulence as a function of observed virulence on seed infected bean plants. Color coding indicates PGs. Dashed line indicates 1:1 relationship. (B) Virulence for each isolate tested ordered by predicted virulence and colored by PG. The solid line represents virulence predictions for each isolate. The grey box represents the error margins for the predictions based on the MAE and RMSE values for the model.

## DISCUSSION

In this work we addressed whether host of isolation is a reliable predictor of host specific virulence and whether whole genome sequences can be used to predict the host specific virulence potential of individual strains. While host of isolation is a widely used surrogate for host specificity, this assumption has rarely been empirically tested [24, 25], and the strength of this assumption is critical when viewed from the context of the virulence potential of emerging pathogens. Are strains isolated from one host only virulent on that host, or do they have the potential to move to other species? Are strains isolated from environmental sources, such as streams or soil, limited to those environments or can they ‘jump’ to a new host and potentially cause a significant outbreak?

Our first aim was to determine if infection of bean seeds by *P. syringae* recapitulated virulence responses seen in standard syringe inoculation virulence assays. We found a negative association between our virulence measure of normalized plant fresh weight after seed infection and *in planta* bacterial growth after syringe infiltration into leaf tissue, showing that the seed infection protocol effectively recapitulates standard methods. This finding is consistent with published and anecdotal reports that infected seed stocks are a significance source of bean disease [45, 46, 51, 52]. In general, we found that bean isolates reduced mean plant fresh weight by 30.2% and median weight by 46.2% compared to non-bean isolates. While the PG2 bean isolates (leaf spot disease caused by pathovar *syringae*) had a normalized mean fresh weight of 0.56±0.293 (SD) compared to 0.64±0.246 for the PG3 bean isolates (halo blight disease caused by pathovar *phaseolicola*), this difference was not significantly different. A similar pattern was found when we examined seed germination frequencies, where bean isolates reduced the average germination frequency by 28.9% and median frequency by 22.0% compared to non-bean isolates, while the 38.8% mean germination frequency of PG2 bean pathogens was significantly lower than the 57.0% mean germination frequency of the PG3 bean pathogens (p=0.014).

We find a striking difference when comparing bean to non-bean isolates found in the same phylogroup. As anticipated, bean seed infection with PG3 bean isolates resulted in significantly higher virulence and lower germination frequency than PG3 non-bean isolates, while in contrast, PG2 bean isolates did not differ significantly from PG2 non-bean isolates. PG2 strains generally (irrespective of host of isolation) show greater virulence on bean, indicating that strains from this phylogroup have lower host specificity, i.e., are host generalists. This is strongly supported when comparing non-bean isolates from PG2 to non-bean PG3 isolates (normalized fresh weight of 0.704 and 0.933, respectively; p= 4.67E-04). These findings are consistent with other studies that have found lower levels of host specificity among PG2 strains [24, 25] and lends support to the hypothesis that PG2 strains may rely as much or more on toxins than T3SEs when compared to other *P. syringae* strains.

We expected that PG3 bean isolates would have higher virulence than PG2 bean isolates since halo blight caused by PG3 pathovar *phaseolicola* is a much more severe disease than spot disease caused by PG2 pathovar *syringae*, but this was not the case. There are several explanations for these data. First, is a simple experimental bias explanation driven by the fact that nearly all the PG3 bean isolates fall into one clonal group as defined by our clonality criteria of a core genome distance of <0.001 average substitutions per site and T3SE profile Jaccard similarity value >0.8. We attempted to address this issue by oversampling from the *phaseolicola* clonal group. But to ensure that we did not create another bias by adding too many very closely related strains, we only added seven additional strains to the original group of six PG3 bean isolates. Unfortunately, this still resulted in a small set that could easily be skewed by a few outlying measurements. Second, some of the PG3 strains likely elicit effector-triggered immunity in the cultivar of bean assayed, which would result in healthy plants. Given the small set of PG3 bean isolates, even a few ETI-eliciting strains will result in a large average decrease in virulence. And third, it is possible that the most severe symptoms of halo blight are only seen after leaf-to-leaf transmission caused by water splash rather than seed transmission [14, 53].

While many genome-wide association studies have successfully identified strong genotype-to-phenotype linkages, we were unable to identify any loci significantly associated with bean isolation (data not shown). Consequently, we shifted our focus to machine learning approaches as they can not only unravel genomic signatures associated with continuous phenotypes, but also predict the virulence potential of previously unseen isolates given their genome sequences. Regardless of PG affiliation, our model was able to predict the virulence of individual *P. syringae* isolates with low error margins based solely on whole genome data. The fact that models trained on virulence factors alone could predict virulence with considerable accuracy supports the notion that T3SEs and phytotoxins play crucial roles in host adaptation processes. Nonetheless, the higher predictive power of models trained with whole genome kmers suggests that factors other than canonically virulence-associated genes also play important roles on disease development and adaptation to beans.

Sample size is one of the most important contributors to accurate model generation in machine learning. Simple models trained with large data sets can learn to predict outcomes with much higher accuracy than complex models trained on smaller datasets. Our results also found a correlation between sample size and model accuracy, with the screened strain collection providing a MAE of 0.153 while the larger imputed strain collection increased model performance to a MAE of 0.065. We also find poorer model performance for PG2 strains than PG3 strains. While this may partly reflect the differences in sample size between these two groups, it also likely reflects the underlying biology. The majority of bean isolates in PG3 are phylogenetically clustered, while there is little clustering of bean isolates in PG2. Consequently, the model may perform better on PG3 since it is essentially predicting phylogenetic structure. Another contributing factor is likely the finding discussed above, namely, that host specificity appears to be weaker in PG2. If PG2 strains are more generalists than specialists, then the host specificity signal would be weaker and any model trying to find this signal would perform more poorly.

## MATERIALS AND METHODS

### Strain collection

Three hundred and thirty-four *Pseudomonas syringae* strains were used in this study (Table S1). Forty-six *P. syringae* isolates were collected from bean fields approximately 80 km East of Lethbridge, Alberta, Canada, during the summer of 2012. Bean leaves with symptoms of bacterial diseases were collected from new growth during the vegetative growth stage. The remaining 288 isolates were previously published [9] and include 49 *P. syringae* type and pathotype strains [1, 35]. A type strain is the isolate to which the scientific name of that organism is formally attached under the rules of prokaryote nomenclature, while a pathotype strain is similar but with the additional requirement that it has the pathogenic characteristics of its pathovar (i.e., a pathogen of a particular host) [5]. Out of the 334, 318 strains were used for comparative analyses and model training, while 16 were used for model functional validation. A subset of 267 non-clonal representative strains (discussed below) were selected for the predictive modeling to avoid clonal bias. A further subset of 121 isolates, including the type and pathotype strains, were selected for virulence assays.

### Sequencing and quality control

DNA was extracted using the Gentra Puregene Yeast and Bacteria kit (Qiagen, Hilden, Germany). Illumina libraries with 300-400 bp inserts were generated using the Illumina Nextera XT kit according to the manufacturer’s protocol (Illumina, CA, USA). Samples were multiplexed with the Illumina Nextera XT Index kit containing 96 indices. Samples were sequenced on the Illumina NextSeq 500 Mid Output v2 (300 cycle) kit with 150 base PE reads. All sequencing was performed at the University of Toronto’s Centre for the Analysis of Genome Evolution and Function (CAGEF). Raw read quality was assessed with FastQC. Trimmomatic was used to remove adapters and trim raw sequencing reads based on a sliding window approach (window size = 4, required quality = 5).

### *De novo* assembly

Paired-end reads were *de novo* assembled using the CLC Genomics Assembly Tool (CLC Genomics Workbench). Contigs shorter than 1kb were removed from the assemblies. Low coverage contigs with matches to non-*Pseudomonas* genera and no matches to the *Pseudomonas* genus that had a depth of coverage less than one standard deviation from the average assembly coverage were deemed contaminants and, therefore, removed from the final draft.

### Pangenome analysis, gene prediction, annotation, and orthologous clustering

Gene prediction and annotation for all assemblies were performed with Prokka [36]. Prokka annotates inferred coding sequences by searching for sequence similarity in the UniProtKB [37] database and HMM libraries [38]. Additionally, all predicted genes were aligned against a custom T3SE database for the identification of potential T3SEs [19]. Pangenome analysis was performed via PIRATE [39], which iteratively clusters genes into orthologous groups by performing all-vs-all comparisons followed by MCL clustering given a certain percent identify threshold. Genes present in at least 95% of the genomes were classified as core. Core protein families were individually aligned with MUSCLE [40] and later concatenated into a single protein alignment. We used the FastTree2 approximate maximum-likelihood approach [41] to infer the phylogenetic relationships of all 320 isolates. Core genome synonymous substitution rates were estimated with MEGA7 (41) using the Nei-Gojobori method and Jukes-Cantor model. Jaccard distances were computed with R version 4.0.5 (42) using a binary matrix of presence and absence of accessory genes. Rarefaction curves were generated using a custom Python script.

### Identification of non-clonal representative strains

We reduced the impact of phylogenetic bias in our predictive modeling by selecting only one representative strain from each clonal group (i.e., very closely related strains recently derived from a common ancestor) identified from the *P. syringae* core-genome phylogeny. We identified clonal groups by calculating the pairwise core genome evolutionary distance and the Jaccard similarity for T3SE profiles. We found the minimum pairwise core-genome evolutionary distance for isolates with identical T3SE profiles to be 0.001 average substitutions per site. We therefore pooled the 318 isolates if they had a core genome evolutionary distance of less than 0.001, resulting in 209 clusters. We further supplemented these clusters by adding back any strain that had a T3SE profile Jaccard similarity value less than 0.8, resulting in 267 non-clonal clusters. A single representative was selected out of each of these non-clonal clusters for downstream analyses. One exception was made to the strain selection process to balance our experimental design, which was skewed due to the fact that the vast majority of PG3 bean isolates (i.e., pathovar *phaseolicola*) fall into one clonal group. Initially, our selection criteria resulted in only six PG3 bean isolates compared to 16 PG2 bean isolates (i.e., pathovar *syringae*). We therefore added an additional seven *phaseolicola* strains to the screened set to better balance the number of bean isolates in PG2 and PG3. Evolutionary distances and Jaccard similarity scores were inferred with MEGA7 [42] and R version 4.0.5 [43].

### Seed infection virulence assay

*P. syringae* strains were grown overnight at 30°C in King’s B media, re-suspended in 10 mM MgSO_4_ and diluted to an OD_600_ of 0.001. *P. vulgaris* var. Canadian Red seeds were soaked for 24 hours in the bacterial suspension, planted in Sunshine Mix 1 soil with regular watering and grown for 14 days. Plant fresh weight and germination frequencies were measured and normalized to a control plant treated with 10 mM MgSO_4_ sown on each flat. Trials were repeated three times.

### Syringe infiltration virulence assays

*P. syringae* strains were grown overnight on appropriate antibiotics, re-suspended in 10 mM MgSO_4_ and diluted to OD_600_ of 0.001. Two- to three-week-old *Phaseolus vulgaris* var. Canadian Red plants were syringe infiltrated and bacterial growth assays were carried out by harvesting eight leaf disks (1 cm^2^) from each plant (two per each primary leaf) three days after infiltration. Disks were homogenized using a bead-beater in 200 μl sterile 10 mM MgSO_4_, serially diluted in 96-well plates, and 5 μl from each dilution was spot plated on KB supplemented with rifampicin for positive and negative control strains, and rifampicin and kanamycin for strains harboring constructs. Plates were incubated for at least 24 hours at 30°C and the resulting colony counts were used to calculate the number of CFUs per cm^2^ in the leaf apoplast.

### Predictive modeling of *P. syringae* virulence on bean

We used an implementation of gradient boosted decision trees to model the effect of *P. syringae* isolates on plant weights as a proxy for strain virulence using: 1) whole genome kmers, 2) T3SE kmers, and 3) a presence / absence matrix of T3SEs and phytotoxins. We split sequences into 31-mers with fsm-lite and generated a binary matrix for kmers with identical distribution patterns using custom python scripts. Next, we used the Scikit-learn and the XGBoost python libraries [44] to generate a regression model for the prediction of normalized plant weights using all three datasets as input features. Given the relatively small size of our dataset, we used a cross-validation (CV) procedure to assess the performance of our model on 50 independent splits. For each time, we randomly split the data into training (80%) and testing (20%) sets while maintaining the same plant weight distributions on both sets. Hyper parameters were fine-tuned using Scikit-learn’s RandomizedSearchCV module and regression models were generated with XGBoost’s XGBRFRegressor module.

## SUPPORTING INFORMATION CAPTIONS

**Figure S1:**
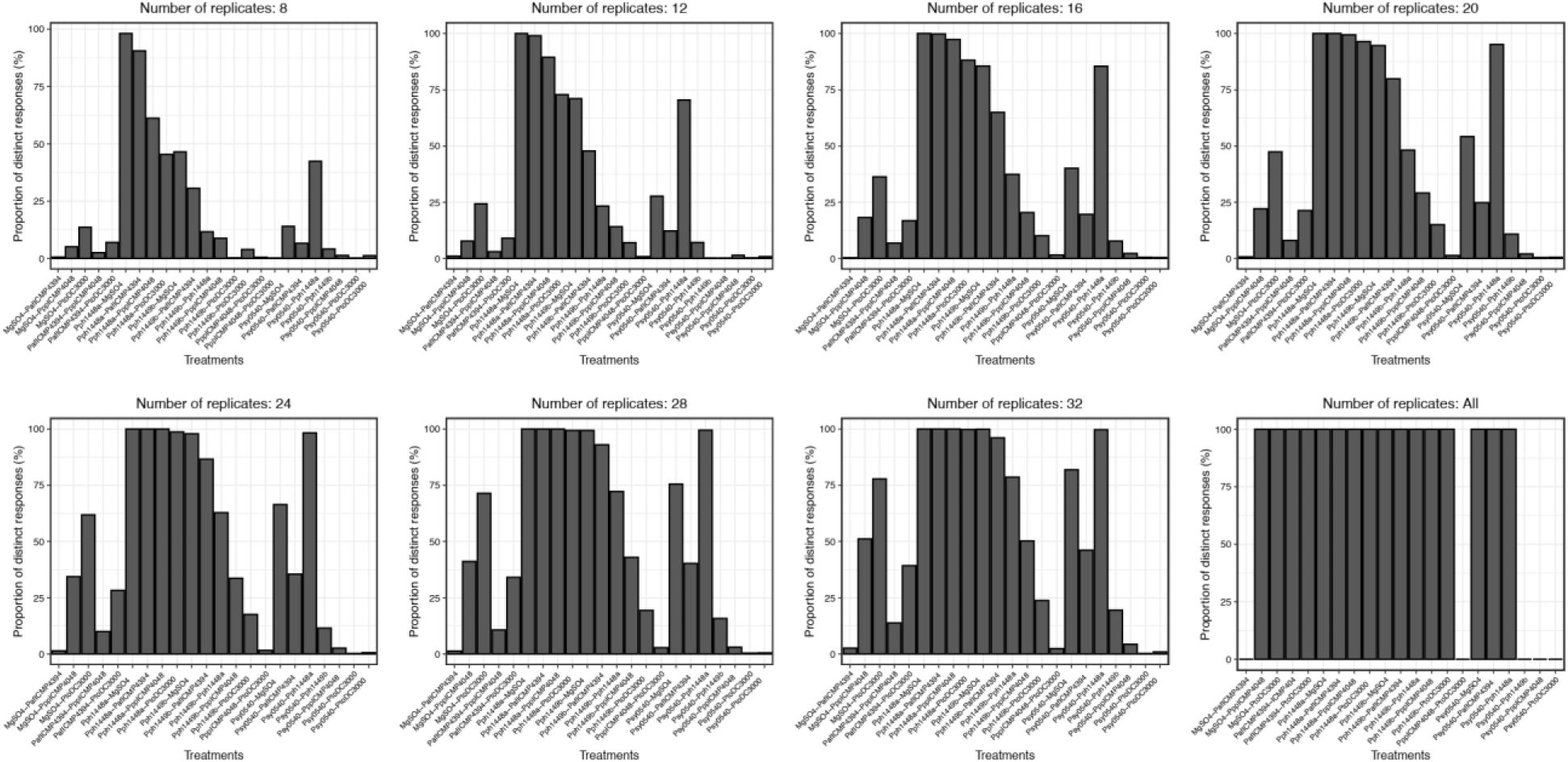
Sample size selection via a simulated experimental setup. Bean plants were seed infected with 6 *P. syringae* isolates with a high number of replicates (>50). Plant weights were randomly selected according to various replicate sizes (8-32). Our ability to distinguish pathogens from non-pathogens using Tukey-HSD tests plateaus at 20 replicates per treatment.

**Figure S2:**
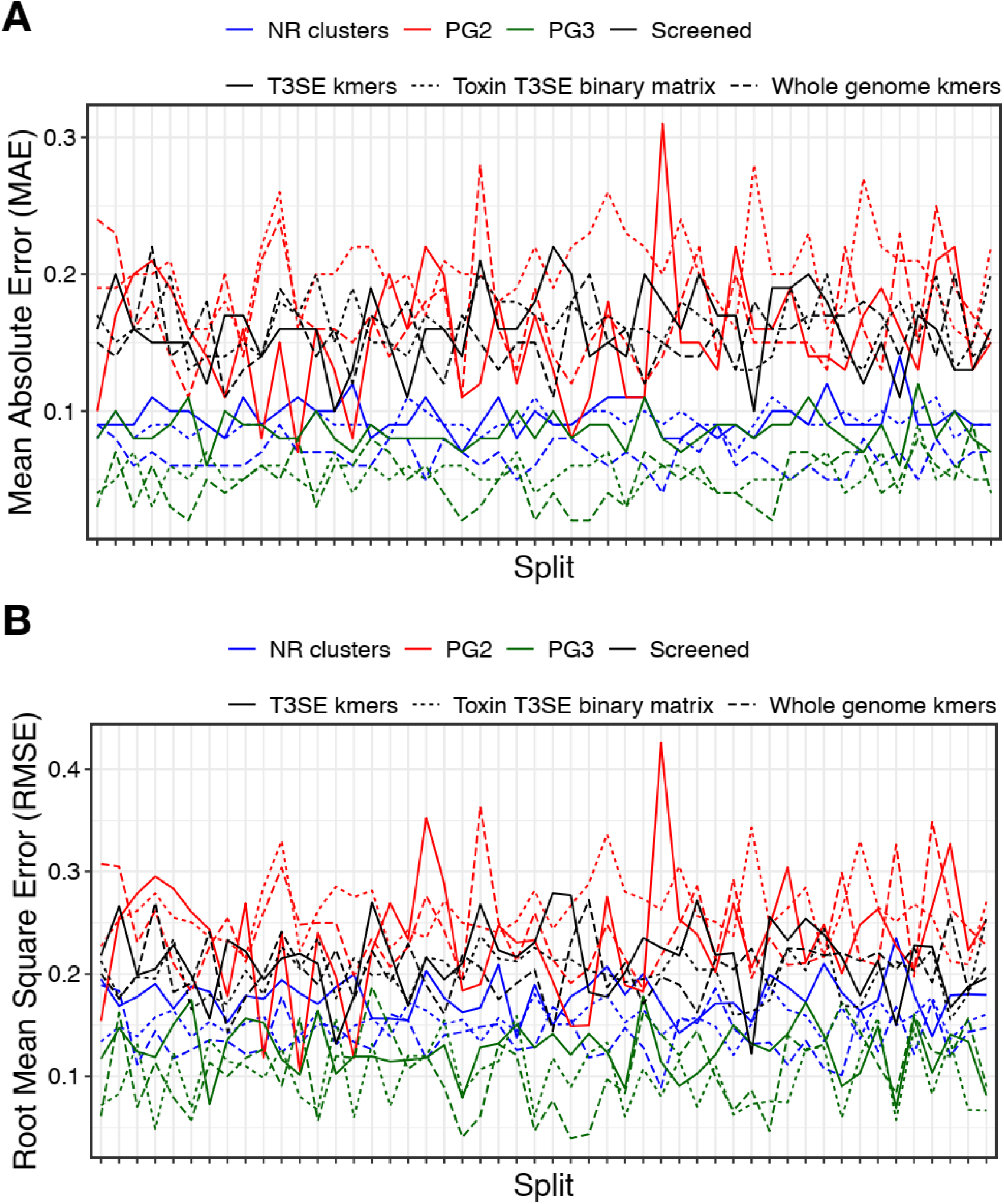
Mean Absolute Error (MAE) and Root Mean Square Error (RMSE) across 50 cross-validation splits. Model performance in terms of (A) MAE and (B) RMSE. Plant weight distributions were kept the same across splits.

**Figure S3:**
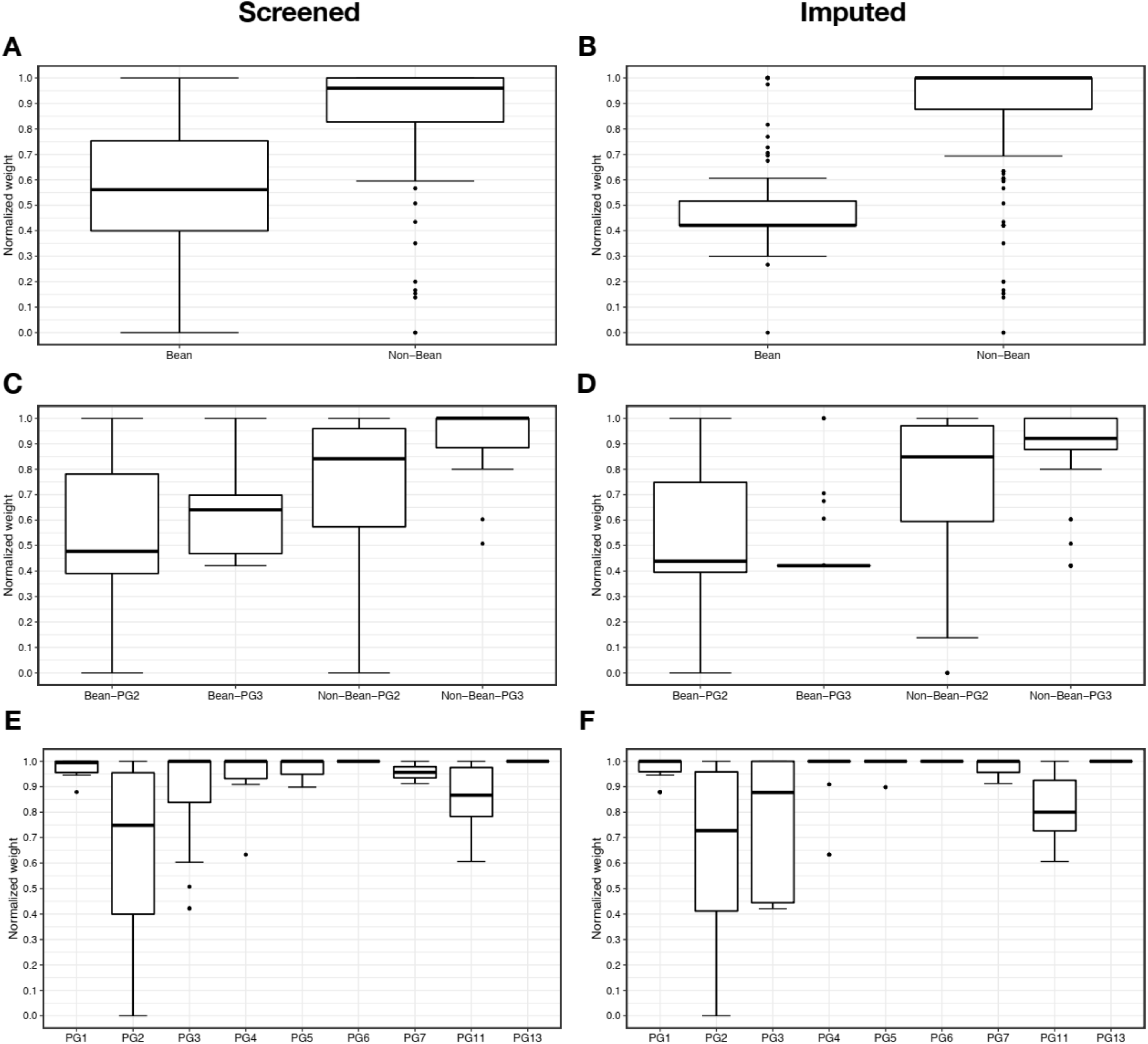
Distribution of virulence values stratified by host and phylogroup. Boxplots showing the distribution of virulence (i.e., normalized plant weight 14 days after seed infection) values for (A) bean verses non-bean isolates for the set of 121 screened strains, and (B) for the 320 strains in the expanded dataset that includes both screened and imputed strains. (C) Distribution of virulence values for bean and non-bean isolates stratified by phylogroup for the screened strains and (D) for the expanded dataset. (E) Distribution of virulence values stratified by phylogroup (PG) for the screened strains, and the (F) expanded strain set.

**Table S1: List of *P. syringae* isolates used in this study.**

